# CEACAM1 Targets ABCA1 to Mediate Atrial Fibrillation via Lipophagy-Dependent Ferroptosis

**DOI:** 10.1101/2024.11.26.625561

**Authors:** Wenwen Hu, Junhua Zou, Zhixuan Li, Boning Yang, Xue Ma, Run Zou, Jing Pu, Ling Zhao, Jing Wang

**Affiliations:** Department of Cardiology, First affiliated Hospital of Kunming Medical University, Kunming, Yunnan Province, China

**Keywords:** Atrial fibrillation, ceacam1, lipophagy, ferroptosis, ABCA1, oxidative stress, gene knockout

## Abstract

Atrial fibrillation (AF) is one of the most common arrhythmias, imposing a significant burden on individuals due to its association with cerebrovascular events. The mechanisms underlying AF involve cardiac structural remodeling, electrical remodeling, and metabolic remodeling, with oxidative stress playing a crucial role. Our previous studies demonstrated that upregulation of CEACAM1 can increase oxidative stress, inhibit cell proliferation, and potentially contribute to cellular damage, while ferroptosis is closely associated with AF. In this study, we observed a significant increase in CEACAM1 expression in an AF model. Knocking out CEACAM1 revealed changes in mitochondrial membrane potential, reactive oxygen species (ROS) levels, intracellular Ca2+ concentration, channel proteins, and glycolipid metabolic enzymes. This knockout reversed electrical, structural, and metabolic remodeling in AF. Further investigation showed that CEACAM1 knockout inhibited ferroptosis, as evidenced by ferroptosis-related indicators. Using colocalization analyses of lipid droplets, lysosomes, and autophagosomes, we discovered that CEACAM1 knockout suppressed autophagy and lipophagy during AF. Supplementation with free fatty acids (FFAs) in the AF model post-CEACAM1 knockout suggested that the inhibition of ferroptosis was linked to a reduction in FFAs. Transcriptomic and non-targeted metabolomic analyses, along with ABCA1 interference studies, indicated that CEACAM1 knockout mitigates AF by promoting ABCA1 expression. In summary, CEACAM1 targets ABCA1 to alleviate AF through the regulation of lipophagy-dependent ferroptosis, as demonstrated in both cellular and animal models.

**Background:** Ferroptosis is a regulated form of cell death, but its connection with atrial fibrillation(AF) remains unclear. Since CEACAM1 knockout reduces oxidative stress and protects cells, we investigated the relationship among CEACAM1, ferroptosis, and AF to uncover the regulatory mechanisms of ferroptosis in AF.

**Methods:** We established AF models using HL-1 cells, human pluripotent stem cells, and rats. CEACAM1 expression levels were assessed in these models, along with changes in mitochondrial membrane potential, ROS levels, intracellular Ca^2+^ concentration, channel proteins, glycolipid metabolic enzymes, and ferroptosis-related indicators following CEACAM1 knockout. Through transcriptomics, non-target metabolomics, and immunoprecipitation screening, we identified ABCA1 as a downstream gene of CEACAM1 involved in protecting against AF. Subsequently, ABCA1 interference experiments were conducted, and colocalization analyses of lipid droplets, lysosomes, and autophagosomes were performed after the application of ferroptosis activators, lipophagy inhibitors, and lipophagy activators. FFAs were added to the AF model to evaluate changes in lipophagy and ferroptosis following CEACAM1 knockout.

**Results:** In the AF model using HL-1 myocytes, CEACAM1 knockout reverse electrical, structural, and metabolic remodeling while inhibiting ferroptosis, with ABCA1 as a downstream gene. In the AF model derived from pluripotent stem cells differentiated into atrial myocytes, CEACAM1 knockout suppressed autophagy and lipophagy during AF, thereby inhibiting ferroptosis, which correlated with reduced FFA release. This process was regulated by ABCA1. In the rat AF model, ABCA1, identified as a downstream gene of CEACAM1, was further confirmed to inhibit ferroptosis by regulating FFA levels.

**Conclusion:** This study reveals that CEACAM1 knockout inhibits ferroptosis in AF models by upregulating ABCA1 and reducing FFA levels, providing a novel regulatory mechanism for ferroptosis in AF.

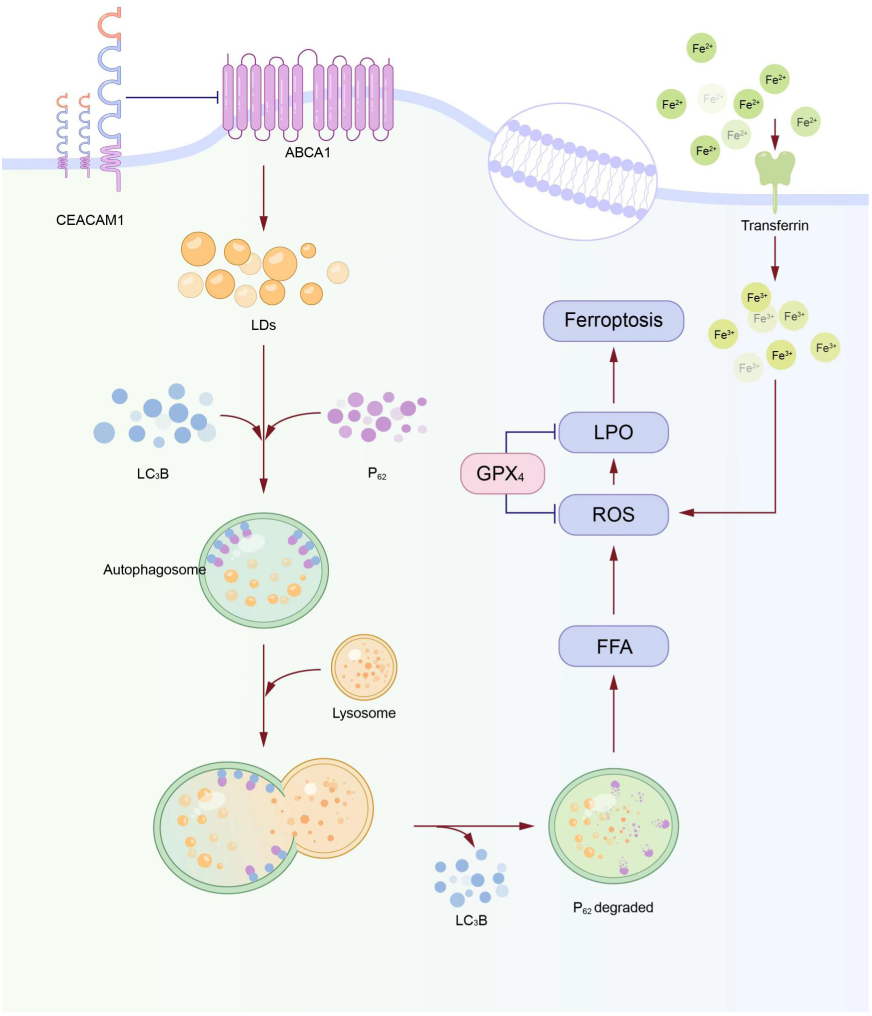

## Introduction

Atrial fibrillation (AF) is one of the most common arrhythmias, with its incidence continuing to rise. The cerebrovascular adverse events caused by AF further exacerbate the burden on human health^1, 2^. The occurrence, development, and persistence of AF are closely related to cardiac structural remodeling, electrical remodeling, metabolic remodeling, and autonomic imbalance. Among these pathological processes, lipid peroxidation, myocardial fibrosis, and inflammation play pivotal roles^3, 4, 5^. Ferroptosis, a recently discovered mode of cell death, is an iron-dependent cascade driven by lipid peroxidation^6, 7^. Studies by Fang et al. revealed that ferroptosis significantly increases susceptibility to AF in a rat model of endotoxemia^8^. Moreover, ferroptosis has been implicated in the progression of hepatic, pulmonary, and renal fibrosis, and its inhibition can prevent tissue fibrosis^9, 10, 11^. Recent findings also suggest that ferroptosis is closely linked to myocardial fibrosis in AF, highlighting its significance in the disease^12^. However, the precise mechanisms and critical mediators driving ferroptosis in AF remain unclear.

CEACAM1 has been identified as an important regulator of immune responses and metabolism, with therapeutic implications in conditions such as cancer and liver injury. Harrson T. et al. found that CEACAM1 in endothelial cells may influence liver fibrosis by reducing endothelin-1 production^13^. Similarly, our prior research demonstrated that upregulation of CEACAM1 in pulmonary vascular endothelial cells under hypoxic conditions increases oxidative stress, inhibits cell proliferation, and induces cellular damage ^14^. Zhang Y.Z. et al. showed that CEACAM1 overexpression in cardiac myocytes infected with coxsackievirus B3 elevated apoptosis and levels of TNF-α and IL-1β^15^. Insights from the 81st Annual Scientific Session of the American Heart Association revealed that Cardiac CEACAM1 plays a central role in regulating cardiac fatty acid metabolism. The loss of cardiac CEACAM1 in Ceacam1−/− mice led to cardiac lipid accumulation and hypertrophy, emphasizing its importance in lipid homeostasis^16^. Najjar S.M. et al. demonstrated that activated CEACAM1 inhibits fatty acid synthase (FAS), thereby reducing fatty acid synthesis and hepatic lipogenesis^17^ . Furthermore, studies by Yun Li and John E. Shively suggested that the interaction between CEACAM1 and β-catenin may regulate Fas-mediated apoptosis in T-cells, fine-tuning the T-cell response^18^. Given these findings, we hypothesize a relationship between CEACAM1, ferroptosis, and AF, potentially involving lipid metabolisim, oxidative stress, apoptosis, and related process. This study aims to further investigate the role of CEACAM1 in atrial metabolic and electrical remodeling in AF, elucidating its role in mediating ferroptosis and its impact on AF pathophysiology.

## Method

### 1 Construction of HL-1 cell atrial fibrillation model

For cells in the logarithmic growth phase, the YC-2-S bipolar programmer was placed in a culture dish and stimulated continuously at 600 times/min and 1V/cm for 24 hours to construct a cell model of atrial fibrillation.

### 2 Detection of protein expression level by WB

After lysing the samples on ice for 10 min, they were centrifuged at 14,000 g for 15 min at 4°C. After measuring the protein concentration with the BCA protein quantification kit, we took 80 μl of protein and 20 μl of 5X protein loading buffer to mix, boiled in a boiling water bath for 5 min, and then performed SDS-PAGE electrophoresis. After the membrane transfer, the PVDF membrane was taken out, and the clipped strips were soaked in 1×TBST and placed in blocking solution (5% skimmed milk powder) to block on a shaker at room temperature for 40 min. Primary antibodies (CEACAM1, ABCA1, Cav1.2, CX40, Kir6.2, LC3, P62, SUR2A) were incubated overnight at 4°C, and secondary antibodies were incubated for 40 min, developed, and photographed for storage.

### 3 CRISPR/Cas9 adenovirus knockout CEACAM1 gene vector construction, Abca1, Fxyd2, Pygm, Cers6 lentiviral interference vector construction and stable cell line construction

The sgRNA sequence of CEACAM1 gene is

TGAAAACTATCGTCGTACTCAGG. Abca1 interfering sequence is

GAGTGCCACTTTCCGAATAAA CTCGAGTTTATTCGGAAAGTGGCACTC TTTTTT, Fxyd2 interfering sequence is

GCAGGIAAIGAAGAIGAACTCHGAGAGTICAICTCATIGACCIGC TTTTT, Pygm interfering sequence is

GAGTTGTACAAGAACCCAAGACTCGAGTCTTGGGTTCTTGTACAAC TC TTTTTT, Cers6 interfering sequence is GCTGACCT

TCACTACTATTACCTCGAGGTAATAGTAGTGAAGGTCAGC TTTTT.

We added 2ul of the plasmid into 100ul of competent Escherichia coli, took a single colony and applied it to 20ml of liquid medium, and cultivated it overnight at 37 degrees Celsius on a 200r shaker. The plasmid was extracted by RT-PCR to screen the best interference vector, and transfected into 293T cells for packaging, and then the cell culture supernatant was collected 48 hours after transfection. Centrifuge at 500g for 10min to remove cell debris for virus titer determination or virus concentration. Transfected cells were continuously selected with 2.5ug/ml puromycin for 1 month.

### 4 Mitochondrial Membrane Potential Detection by Flow Cytometry

We prepared the cells into a single cell suspension, centrifuged at 1000 rpm for 5 minutes, added 0.5ml JC-1 staining working solution, and incubated at 37°C for 20 minutes. Centrifuge at 600g for 3-4 minutes at 4°C to pellet the cells. After washing with JC-1 staining buffer (1X) and resuspending, the cells were analyzed by flow cytometry.

### 5 Detection of reactive oxygen species by flow cytometry

We collected the cells and grouped and labeled them and centrifuged at 1300rpm for 5min. After that, we added 1 ul of DCFH-DA to 1 ml of serum-free medium to resuspend the cells, and incubated in a cell culture incubator at 37°C for 20 minutes before adding 1 ul of Rosup positive Incubate again for 10 min. Add 400μL of serum-free culture medium, mix well, and put it on the machine immediately.

### 6 Immunofluorescence experiment

We fixed cells by incubating in 4% PFA solution for 30 minutes at room temperature, then blocked samples for 1 hour in PBS containing 2% Triton X-100 (PBST) and 5% goat serum, and incubated them with the following primary antibodies Incubate overnight at 4°C: LC3B, NR2F2, α-actinin, LAMP2, PLIN2. We then incubated the cells with the secondary antibody for 1 hr at 37°C. We then imaged and analyzed the cells under a fluorescence microscope.

### 7 Colocalization of lipophage and lysosomes detected by immunofluorescence

We remove the cell culture medium, add Lyso-Tracker Red staining working solution pre-incubated at 37°C, and incubate for 50-60 minutes. Then we removed the Lyso-Tracker Red staining solution; we added 0.5ml 2×BODIPY493/503 into 0.5ml cells and incubated for 30min. Fresh cell culture fluid is then added, followed by observation, usually with a fluorescence or confocal microscope.

### 8 ELISA to detect the levels of PK, LDH, PDH, PDK, GSK-3β, FAT/CD36, ACAD, etc

We collected samples and performed enzyme-linked immunosorbent assays according to the manufacturer’s protocol, including PK, LDH, PDH, PDK, GSK-3 β, FAT/CD36, ACAD, MDA, LOP, ferrous ion, NADPH, GSH, and GSSG, TG and FFA.

### 9 Transcriptome combined with non-target metabolome sequencing

We carried out total RNA extraction and quality inspection, construction and evaluation of sequence analysis library, RNA-seq sample sequencing and screening, reference sequence comparison analysis, gene expression level analysis, gene expression difference analysis and a series of data processing and screening on six replicate cell samples in Control group, AF group and sgCEACAM1+AF group. Finally, differentially expressed genes and significantly enriched signaling pathways were screened out.

### 10 RT-PCR to verify the expression of differential genes

We extracted the total RNA of each group separately, then performed reverse transcription reaction, and then carried out qPCR reaction with 2 x Universal Blue SYBR Green qPCR Master Mix kit. The reaction program was: 95°C pre-denaturation for 1 min; 95°C denaturation for 20s, 55°C annealing 20s, 72°C extension for 30s; 40 cycles. Primer sequences are shown in Table 1.

### 11 Co-immunoprecipitation detection of the interaction relationship between CEACAM1 and Pygm and ABCA1

We took 30ul protein supernatant to prepare total protein (input) and denatured it in boiling water for 10min. Then we added 10ul magnetic beads to 3ug protein and incubated on a shaker at room temperature for 4h/4° overnight, one of which was added with CEACAM1 (pygm) antibody, labeled as IP, and one with HRP-labeled goat anti-rabbit antibody labeled as IgG. After mixing them, they were incubated overnight at 4°C, and rewarmed at room temperature for 1 hour, then we used magnetic poles to absorb magnetic beads, discarded the supernatant, and denatured them in boiling water for 10 minutes. Then we carried out SDS-PAGE electrophoresis, and after the electrophoresis, the gel was taken for silver staining.

### 12 Differentiation of human pluripotent stem cells into human atrial myocytes

We added human pluripotent stem cells in the logarithmic growth phase to 2mL CardioEasy® Human Cardiomyocyte Differentiation Complete Medium I, II and III respectively, and cultured them in a CO2 incubator at 37°C for 48 hours at intervals. On the 11th to 15th day, we used lactic acid screening to obtain relatively pure cardiomyocytes. On the 15th to 20th day, we gave 1 μM retinoic acid treatment, and changed the medium every day until the 25th to 30th day, and collected the cells for subsequent detection.

### 13 Atrial Fibrillation Animal Model Construction

We injected AAV/CAS9-CEACAM1 1x1012GC/ml (200ul) into the tail vein of 350±25g 18-week-old SPF male SD rats for one week of adaptive feeding. Two weeks later, we injected 0.1 mL of Ach-CaCl2 mixture (containing Ach 33 μg/mL + CaCl2 5 mg/mL) into the tail vein again for 7 consecutive days to establish the AF model. Then we inserted the metal needle under the skin of the sterilized extremities, dipped the end of the lead in saline and clipped it to the exposed end of the needle, connected it to a Power Lab 15T multi-conductivity physiological recorder, measured the electrocardiogram of lead II, and observed it dynamically for 5 minutes. The second lead electrocardiograms of the rats were detected before and after modeling.

### 14 HE and Masson staining

We fixed atrial tissue overnight in 4% paraformaldehyde, dehydrated it through an ethanol gradient, embedded it in paraffin, and sectioned it at 5 μm thickness. After staining with hematoxylin and eosin and Masson’s trichrome, the slides were sealed, and a randomly selected area was photographed and observed with an optical microscope.

### 15 Observation of mitochondrial changes by transmission electron microscope

We took 1 mm3 of fresh heart tissue and fixed it in electron microscope fixative: 1% osmic acid in the dark and then fixed at room temperature for 2 hours; dehydration, osmotic embedding, polymerization and sectioning were performed successively, and stained in 2% uranyl acetate saturated alcohol solution in the dark 8min; then stained in 2.6% lead citrate solution to avoid carbon dioxide for 8min; finally dried overnight at room temperature, we observed under a transmission electron microscope and collected images for analysis.

### 16 Detection of intracellular Ca2+ by flow cytometry

We took an appropriate amount of cells to be detected and added them to the Fluo-4 AM working solution, and incubated at 20°C-37°C for 10-60 minutes before loading the fluorescent probes, then washed them with PBS, and tested them on a flow cytometer.

### 17 Statistical Analysis

We used Prism graphpad for statistical analysis, and the experimental data were expressed in the form of mean ± standard error (Mean ± SEM). The comparison of measurement data between groups was performed by T test, and the comparison of measurement data among multiple groups was performed by one-way analysis of variance. Set p < 0.05 to be statistically significant: p < 0.01 to be statistically significant; p < 0.001 to be very statistically significant.

## Results

### 1. Impact of CEACAM1 Knockout on Metabolic Remodeling, Electrical Remodeling, and Ferroptosis in Atrial Fibrillation Myocytes

The correlation between CEACAM1 and AF is illustrated in Figure S1. AF induces both electrical and metabolic remodeling in the atrium. To investigate the role of CEACAM1 in atrial metabolic and electrical remodeling during AF, we analyzed mitochondrial membrane potential, reactive oxygen levels, intracellular Ca2+ concentration, channel protein expression, and glycolipid metabolism enzyme activity following CEACAM1 knockout in HL-1 cell model of AF. Compared with the AF model, CEACAM1 knockout resulted in a significant increase in the percentage of JC-1 (Figures S2 A and S2B), a reduction in ROS levels (Figure S2 C), and a decrease in Ca^2+^ concentration (Figure S2 D). Protein expression levels of Cav 1.2 and CX40 were elevated, whereas Kir6.2 and SUR2A expression levels were reduced (Figures S2 E and S2 F). Key enzymes involved in carbohydrate and lipid metabolism exhibited distinct changes: the expression of PK, LDH, FAT/ CD36, and GSK-3 β decreased significantly, while PDH, PDK, ACAD, CPT-1, and PPARα expression levels increased (Figure S2 G).

Ferroptosis-related indicators were also evaluated after CEACAM1 knockout. The levels of lipid peroxidation (LPO), malondialdehyde (MDA), total iron (Fe), and ferrous iron (Fe^2+^) were all significantly reduced, while NADPH and the GSH/GSSG ratio were increased (Figure 1A). However, when Erastin, a ferroptosis activator, was added to the CEACAM1 knockout model, ROS levels increased (Figure 1B), the percentage of JC-1 decreased (Figure 1C), and intracellular Ca^2+^ concentration was elevated (Figure 1D). Correspondingly, the protein expression levels of Cav1.2 and CX40 were reduced, while Kir6.2 and SUR2A levels increased (Figure 1E). Additionally, LPO, MDA, Fe, and Fe2+ levels were elevated, and the GSH/GSSG ratio and NADPH levels decreased (Figure 1F). These findings indicate that CEACAM1 is closely associated with ferroptosis. Its knockout can significantly inhibit ferroptosis in AF models, suggesting a protective role for CEACAM1 deficiency in mitigating ferroptosis-induced damage.

**Figure 1.**
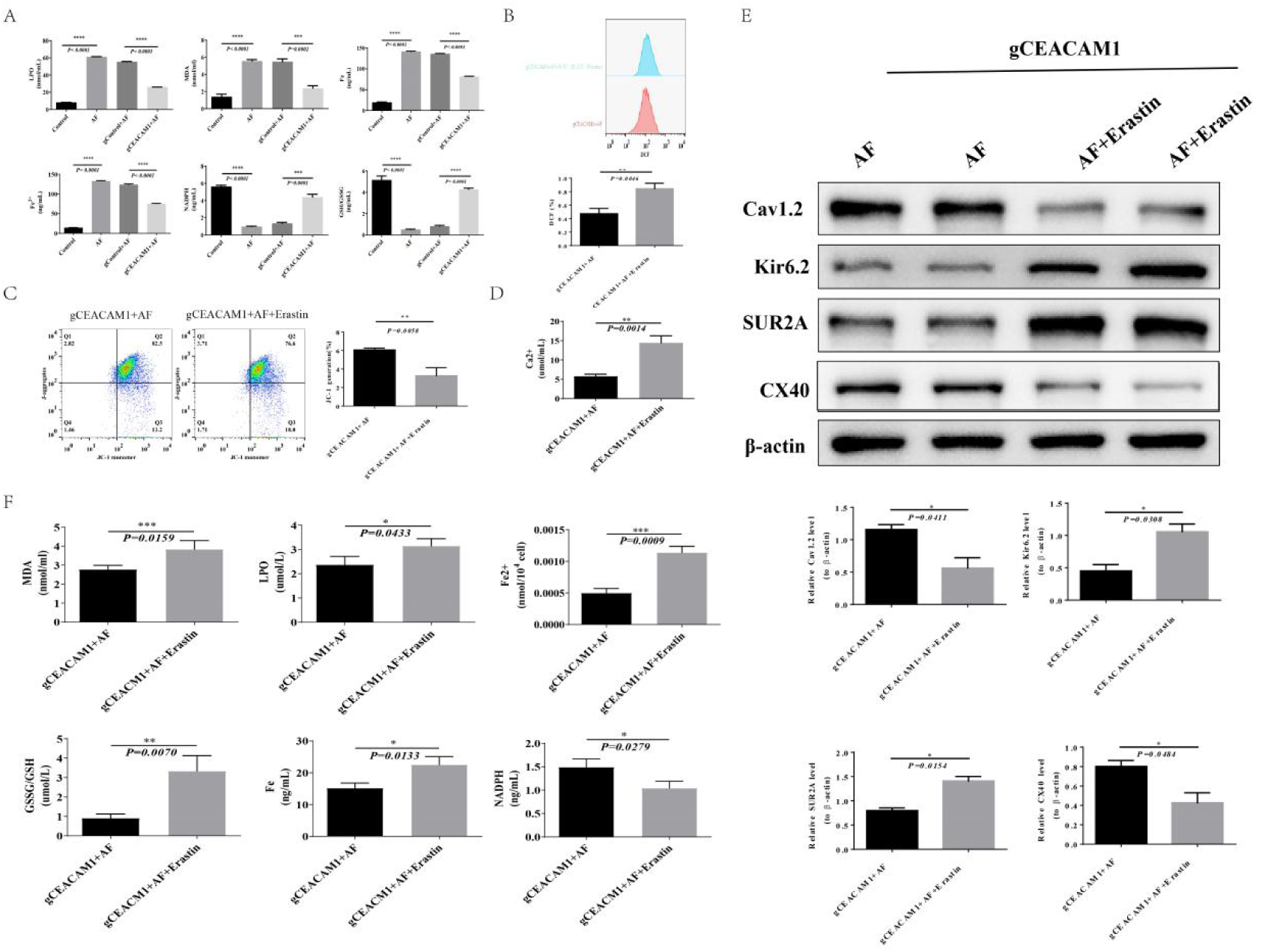
Knocking out of CEACAM1 inhibits ferroptosis in atrial myocytes. A, ELISA analysis showed that LPO, MDA, Fe, and Fe^2+^ levels decreased, while NADPH and GSH/GSSG levels increased after CEACAM1 knockout. B,Flow cytometry demonstrated increased ROS levels after the addition of Erastin to the CEACAM1 knockout model. C, Flow cytometry revealed a reduction in the percentage of JC-1 after Erastin treatment; D Intracellular Ca^2+^ concentrations were elevated after Erastin treatment.; E Western blot analysis indicated changes in channel protein expression: Cav1.2 and CX40 levels decreased, while Kir6.2 and SUR2A levels increased following Erastin treatment. F ELISA results showed increased LPO, MDA, Fe, and Fe^2+^ levels and reduced GSH/GSSG and NADPH levels after adding Erastin.

### 2. Identifying Key Downstream Genes of CEACAM1 That Influence Ferroptosis in Atrial Fibrillation

We identified 119 differentially expressed genes (DEGs) between the two groups (Figure S3A). A clustering heatmap of these DEGs is shown in Figure S3B, a GO enrichment histogram in Figure S3C, and a KEGG pathway enrichment scatter plot in Figure S3D. A total of 80 signaling pathways were enriched in the AF and Control groups (the top 20 pathways with the highest significance are displayed), while 174 pathways were enriched in the AF and sgCEACAM1+AF groups (the top 20 are shown). In the metabolomic analysis, 30 differential metabolites were identified between the AF and Control groups, including 9 up-regulated and 21 down-regulated metabolites. Between the AF and sgCEACAM1+AF groups, 37 differential metabolites were identified, with 24 upregulated and 13 downregulated. Among these, 22 metabolites overlapped between the two comparisons (Figure S3E). KEGG pathway analysis revealed that 35 pathways were enriched between the AF and Control groups, of which 13 showed significant differences. Similarly, 32 pathways were enriched between the AF and sgCEACAM1+AF groups, with 20 pathways displaying significant differences (Figure S3F).

To prioritize key genes and metabolites, we used Venn diagram intersection analysis. No shared pathways were enriched between the transcriptomic and metabolomic analyses for the AF vs. Control comparison. However, for the AF vs. sgCEACAM1 comparison, 9 pathways were jointly enriched in differential genes and metabolites (Figure S4A). The expression levels of DEGs are shown in Figure S4B. Among the key metabolites, triglycerides (TG) and cholesterol esters (CE) were significantly downregulated in the sgCEACAM1+AF group (Figure S4C). Differential expression trends of Abca1, Fxyd2, Pygm, and Cers6 were opposite to those observed for other DEGs (Figure S4D). Consequently, Abca1, Fxyd2, Pygm, and Cers6 were selected for further investigation.

The construction of stable interference strains for Abca1, Fxyd2, Pygm, and Cers6 is illustrated in Figure S5. Subsequent analyses revealed that JC-1 fluorescence significantly decreased in the sh-Pygm group compared to the AF group (Figure S6A). Western blotting confirmed the interference efficiency for Abca1, Fxyd2, Pygm, and Cers6 (Figure S6B). ROS levels increased in the sh-Abca1 and sh-Pygm groups (Figure S6C), as did intracellular Ca^2+^ concentrations (Figure S6D). Channel protein expression levels also varied: Cav1.2, SUR2A, and Kir6.2 increased in the sh-Abca1 and sh-Pygm groups, while CX40 decreased. In contrast, SUR2A and Kir6.2 increased in the sh-Cers6 group, and CX40 increased in the sh-Fxyd2 group (Figures S6E and S6F). Metabolic enzymes showed distinct trends across groups. In the sh-Abca1 group, PK, LDH, FAT/CD36, and GSK-3β levels significantly increased, while PDH, PDK, ACAD, CPT-1, and PPARα decreased. In the sh-Fxyd2 group, PPARα levels increased, but PK, PDH, PDK, GSK-3β, and FAT/CD36 levels decreased. The sh-Pygm group exhibited increases in PK, LDH, FAT/CD36, and GSK-3β, but decreases in PDH, PDK, ACAD, CPT-1, and PPARα. In the sh-Cers6 group, LDH and PPARα increased, whereas GSK-3β decreased (Figure S6G). These findings suggest that sh-Abca1 and sh-Pygm are the two most critical downstream genes associated with atrial metabolic and electrical remodeling.

To further explore how CEACAM1 downstream genes affect ferroptosis in atrial myocytes, we analyzed ferroptosis-related factors and key metabolites using ELISA (Figure 2). Compared to the sh-NC group, LPO, MDA, Fe, Fe^2+^, TG, and CE levels increased in the sh-Abca1 and sh-Pygm groups, while NADPH and GSH/GSSG levels decreased. In the sh-Fxyd2 and sh-Cers6 groups, NADPH levels also decreased; however, LPO levels increased in the sh-Fxyd2 group, while TG and CE levels decreased in the sh-Cers6 group. These results suggest that sh-Abca1 and sh-Pygm are significantly implicated in ferroptosis.

**Figure 2.**
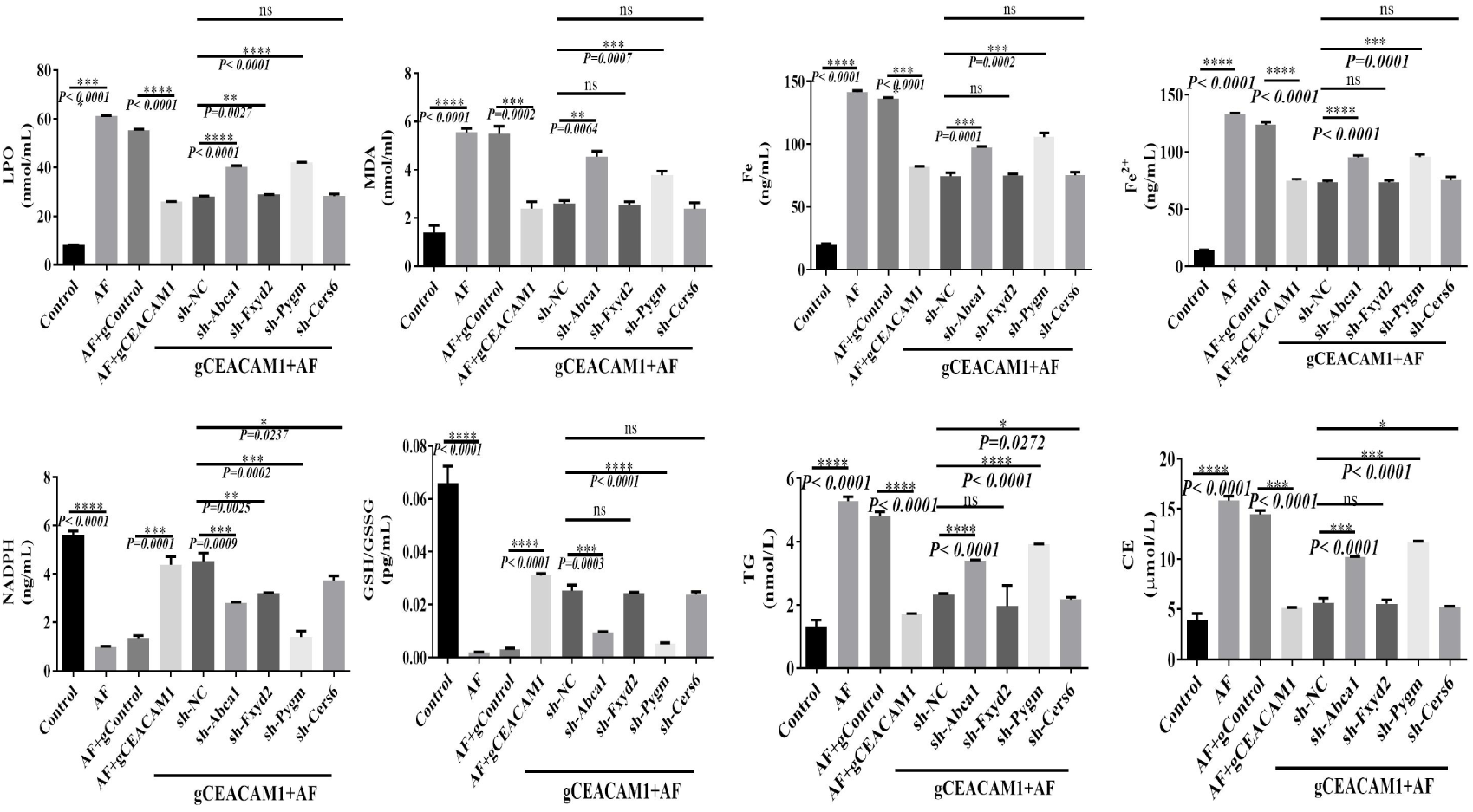
CEACAM1 Knockout Reveals Key Downstream Genes Affecting Ferroptosis in Atrial Myocytes.

Co-immunoprecipitation was performed to identify proteins interacting with CEACAM1 (Figure S7 A). Among these, only ABCA1 was pulled down in the immunoprecipitated samples (Figure S7 B). Cellular and organelle morphology was further observed under electron microscopy. In the sh-Abca1 group, cells showed intact membranes, complete nuclear membranes, and uniform chromatin distribution. Mitochondria were slightly swollen with partially broken cristae, and rough endoplasmic reticulum (RER) exhibited mild expansion with some ribosomal degranulation. No lipid droplets or autophagosomes were observed, but 31 autophagolysosomes (ASS) were noted. In the sh-Pygm group, cells exhibited slight edema and partial cell membrane damage (black arrow). Nuclei were irregular but maintained intact membranes, with evenly distributed chromatin. Mitochondria were slightly swollen with a small number of intact cristae. RER showed moderate expansion and mild ribosomal degranulation. Numerous lipid droplets were present, but no autophagosomes were observed, with 21 autophagolysosomes detected (Figure S7 C). Based on these findings, ABCA1 was selected as the key downstream gene of CEACAM1 for subsequent studies.

### 3. Mechanism of CEACAM1 Downregulation of ABCA1 and Its Induction of Lipophagy and Ferroptosis in Human Atrial Myocytes

We constructed human pluripotent stem cells with CEACAM1 knockout (Figure S8) to study the effect of CEACAM1 on lipophagy during atrial fibrillation (AF). Co-localization analysis of lipid droplets, lysosomes, and autophagosomes revealed significantly reduced co-localization after CEACAM1 knockout (Figures S9 A, S9 B). Protein analysis showed an increase in P62 expression and a decrease in LC3BII levels (Figure S9C), accompanied by reductions in free fatty acids (FFA) and triglycerides (TG) (Figure S9 D). These results demonstrate that CEACAM1 knockout inhibits autophagy and lipophagy during AF, particularly affecting FFA release.

Next, we applied the lipophagy inhibitor LAL and the lipophagy inducer Rapa to assess their effects on lipid droplet localization. In the AF+LAL group, lipid droplet co-localization with lysosomes and autophagosomes was significantly reduced compared with the AF group, while co-localization increased in the AF+g CEACAM 1+Rapa group compared with the AF+g CEACAM1 group (Figure 3A). TG increased and FFA decreased in the AF+LAL group relative to the AF group; both TG and FFA increased in the AF+gCEACAM 1+Rapa group relative to the AF+gCEACAM1 group (Figure 3B). DCF percentages, reflecting reactive oxygen species (ROS) levels, decreased in the AF+LAL group relative to the AF group, while they increased in the AF+gCEACAM1+Rapa group relative to the AF+gCEACAM1 group (Figure 3C). In ferroptosis-related indicators, LPO, MDA, Fe, and Fe^2+^ levels decreased in the AF+LAL group compared with the AF group, while NADPH levels increased. Conversely, these markers increased, and NADPH decreased in the AF+gCEACAM1+Rapa group compared with the AF+gCEACAM1 group (Figure 3D). Regarding autophagy-related proteins, LC3BII levels decreased and P62 levels increased in the AF+LAL group relative to the AF group. In contrast, LC3BII increased and P62 decreased in the AF+gCEACAM 1+Rapa group relative to the AF+gCEACAM1 group (Figure 3E). These findings suggest that CEACAM1 may regulate lipophagy through ABCA1 modulation.

**Figure 3.**
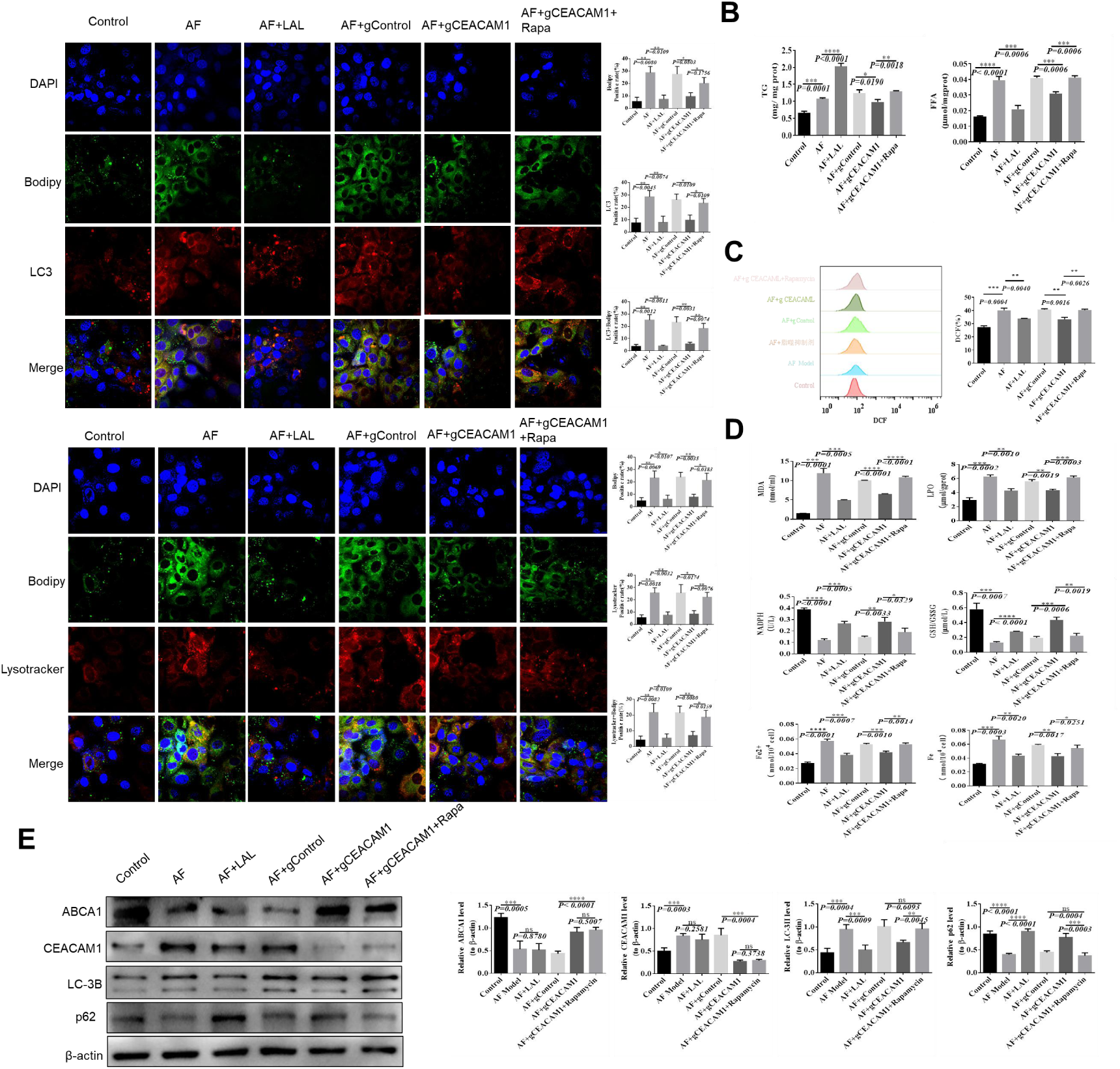
Knocking out of CEACAM1 prevents ferroptosis by inhibiting lipophagy in atrial myocytes in AF. A, Co-localization of lipid droplets with lysosomes and autophagosomes. B, Changes in TG and FFA content. C, Detection of intracellular ROS. D, Measurement of ferroptosis-related factors. E Analysis of autophagy-related proteins.

### 4. Knocking Out CEACAM1 Inhibits Ferroptosis by Reducing FFA Levels

To determine whether CEACAM1 knockout inhibits ferroptosis through FFA reduction, we added FFA (2mM) to the AF model with CEACAM1 knockout and conducted tests after 24 hours Compared with the gControl+AF group, the gCEACAM1+AF group showed reduced co-localization of lipid droplets with lysosomes (Figure 4A) and decreased TG and FFA levels (Figure 4B). The DCF percentage, indicating ROS levels, was also reduced (Figure 4C). Additionally, ferroptosis markers LPO, MDA, Fe, Fe2+, GSH/GSSG decreased, while NADPH levels rose (Figure 4D). In contrast, adding FFA to the gCEACAM1+AF group (gCEACAM1+AF+FFA) reversed these effects. Co-localization of lipid droplets and lysosomes increased (Figure 4A), TG and FFA levels rose (Figure 4B), and the DCF percentage increased (Figure 4C). Ferroptosis-related markers, including LPO, MDA, Fe, Fe2+, GSH/GSSG also increased, while NADPH decreased (Figure 4D). These results indicate that CEACAM1 knockout suppresses ferroptosis by reducing FFA content.

**Figure 4.**
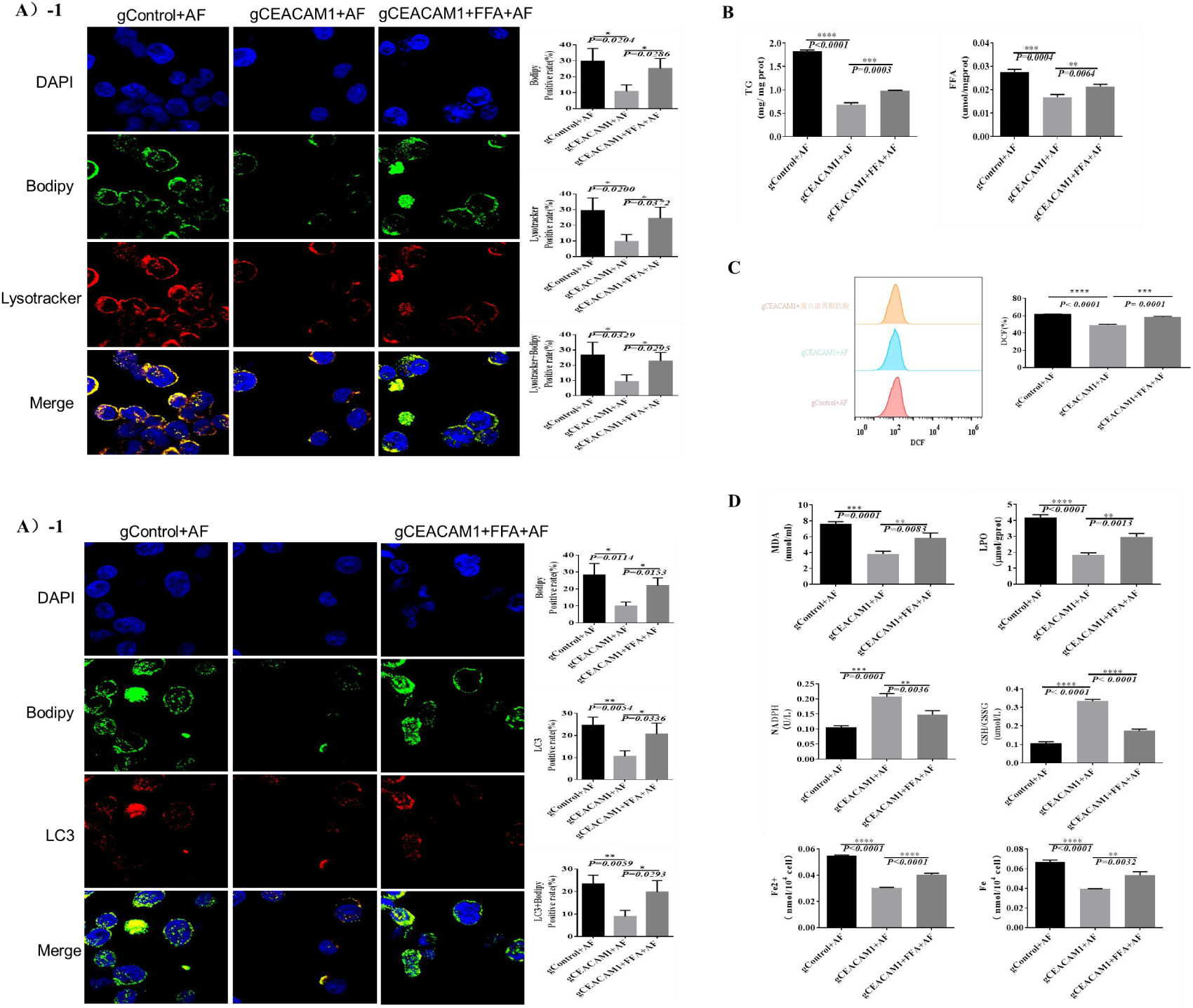
Knocking out of CEACAM1 inhibits ferroptosis by reducing FFA content. A,Co-localization of lipid droplets with lysosomes and autophagosomes; B,Changes in TG and FFA levels; C, Detection of intracellular ROS; D,Analysis of ferroptosis-related factors.

### 5. Knocking Out CEACAM1 Upregulates ABCA1, Inhibiting Lipophagy to Reduce FFA Levels and Suppress Ferroptosis

To investigate whether the up-regulation of ABCA1 following CEACAM1 knockout inhibits lipophagy and reduces FFA content to suppress ferroptosis, we performed ABCA1 interference in CEACAM1 knockout cells. The results showed that, compared with gCEACAM1+AF group, the gCEACAM1+sh-Abca1+AF group exhibited increased levels of FFA and TG (Figure 5C), an elevated DCF percentage (Figure 5D), and increased levels of LPO, MDA, Fe, Fe^2+^, and GSH/GSSG, while NADPH decreased (Figure 5E). Additionally, LC3BII levels were elevated, P62 levels decreased, and CEACAM1 levels remained unchanged (Figure 5F). Mitochondrial changes were also observed. In the gCEACAM1+AF group, mitochondria (M) showed mild swelling, slightly lighter matrices, a small number of intact cristae, and occasional breaks, with no apparent lipid droplets or autophagosomes detected. In contrast, mitochondria in the gCEACAM1+sh-Abca1+AF group exhibited mild swelling with uneven matrices, a few intact cristae that were sometimes shorter or broken, an intact membrane structure, and increased lipid droplets (LD) within the cells, but no noticeable autophagosomes (Figure 5G). These findings indicate that ABCA1, as a downstream gene of CEACAM1, plays a role in regulating FFA content and inhibiting ferroptosis.

**Figure 5.**
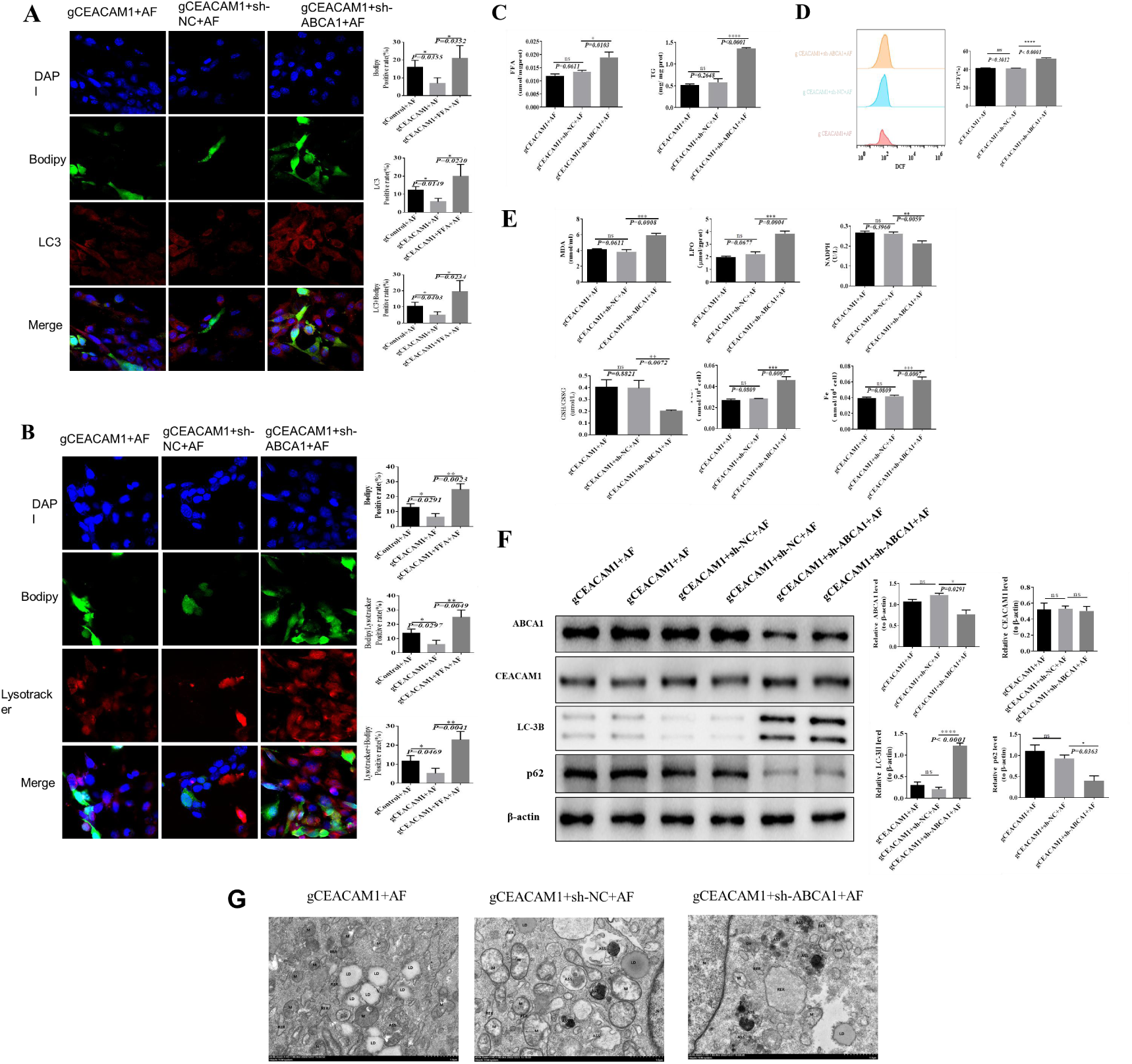
Knocking out CEACAM1 inhibits ferroptosis by upregulating ABCA1, which suppresses lipophagy and reduces FFA content. A, Co-localization of lipid droplets and lysosomes; B, Co-localization of lipid droplets and autophagosomes; C,Detection of TG and FFA content changes; D, Detection of intracellular reactive oxygen species; E, Detection of ferroptosis-related factors; F, Detection of autophagy-related proteins; G, Electron microscopy imaging of mitochondrial changes.

### 6. Protective Effect of CEACAM1 Knockdown on Atrial Fibrillation In Vivo

An atrial fibrillation (AF) model was established in SD rats. Electrocardiogram results confirmed successful model establishment, as evidenced by the disappearance of the P wave and irregular P-R intervals in the model group (Figure 6A). HE staining showed that CEACAM1 knockout significantly alleviated myocardial injury (Figure 6B), reduced myocardial fibrosis (Figure 6C and 6D), and increased Abca1 protein and mRNA expression levels (Figure 6E and 6F). These results suggest that knocking down CEACAM1 provides protection against AF, likely via the promotion of ABCA1 expression.

**Figure 6.**
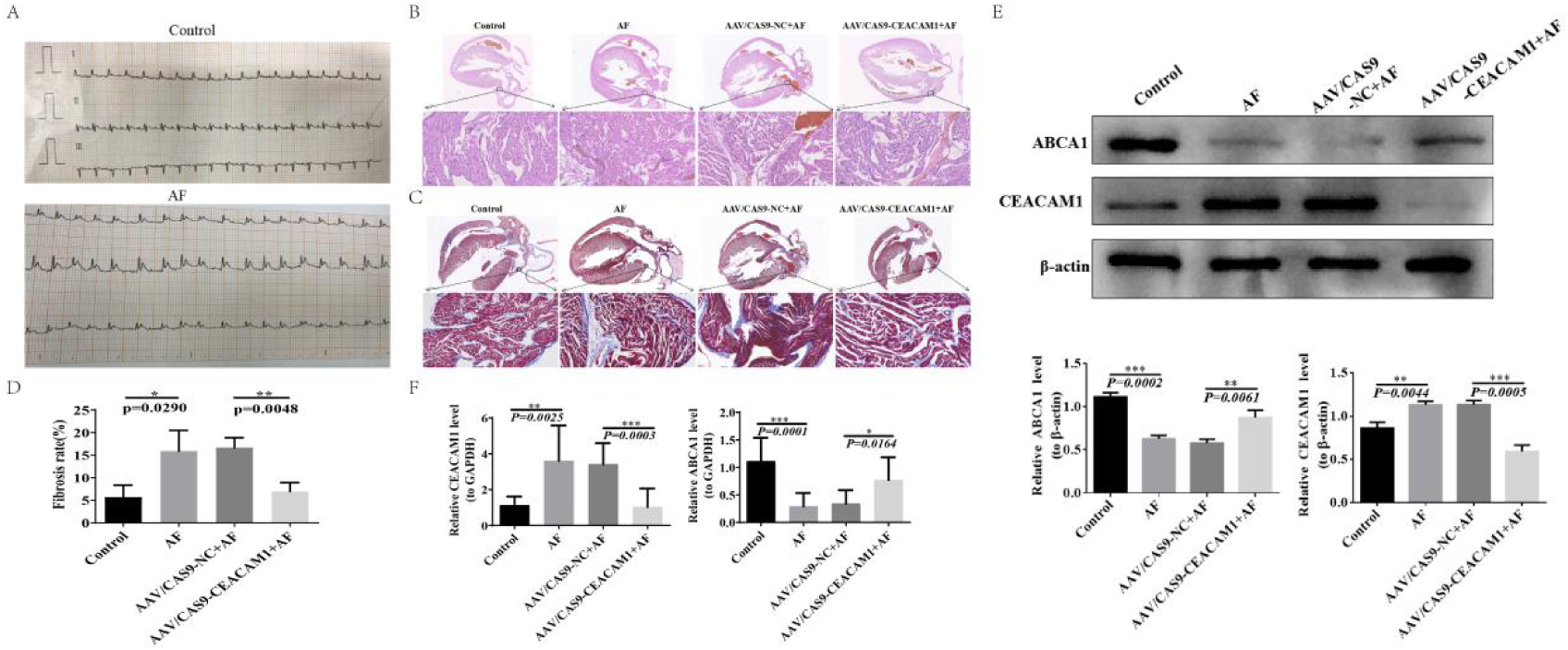
In vivo study of the protective effect of CEACAM1 knockdown in atrial fibrillation. A, Electrocardiogram results; B, HE staining; C, Masson staining; D, Quantification of fibrosis by Masson staining; E, Changes in CEACAM1 and ABCA1 protein expression; F, Changes in CEACAM1 and ABCA1 mRNA expression.

### 7. Mechanism of CEACAM1 in Atrial Fibrillation: Inducing Lipophagy, Releasing FFA, and Promoting Ferroptosis in Atrial Myocytes

We investigated TG and FFA levels, mitochondrial morphology, autophagy-related proteins, ferroptosis-related factors, and key proteins in lipid metabolism. CEACAM1 knockout reduced TG and FFA levels (Figure 7A), increased P62 expression, and decreased LC3BII levels (Figure 7C). Furthermore, levels of LPO, MDA, Fe2+, and GSH/GSSG decreased, while NADPH levels increased (Figure 7D). FAT/CD36 expression decreased, whereas CPT1 and PPARα levels increased (Figure 7E). Mitochondrial observations revealed differences between groups. In the Control group, most of the mitochondria (M) were intact, with no swelling, uniform matrices, clear cristae, and occasional membrane protrusion or blurring. In the AF group, most mitochondria were severely swollen, enlarged in volume, with dissolved matrices, broken cristae, and local vacuolization. Following CEACAM1 knockout, mitochondrial abundance increased, with most showing no swelling and uniform matrices. A small proportion displayed slight swelling, uneven matrices, and tolerable structural damage, such as shortened or broken cristae (Figure 7B). These findings align with in vitro results, indicating that that CEACAM1 induces AF by promoting lipophagy in atrial myocytes, thereby releasing FFA and triggering ferroptosis.

**Figure 7.**
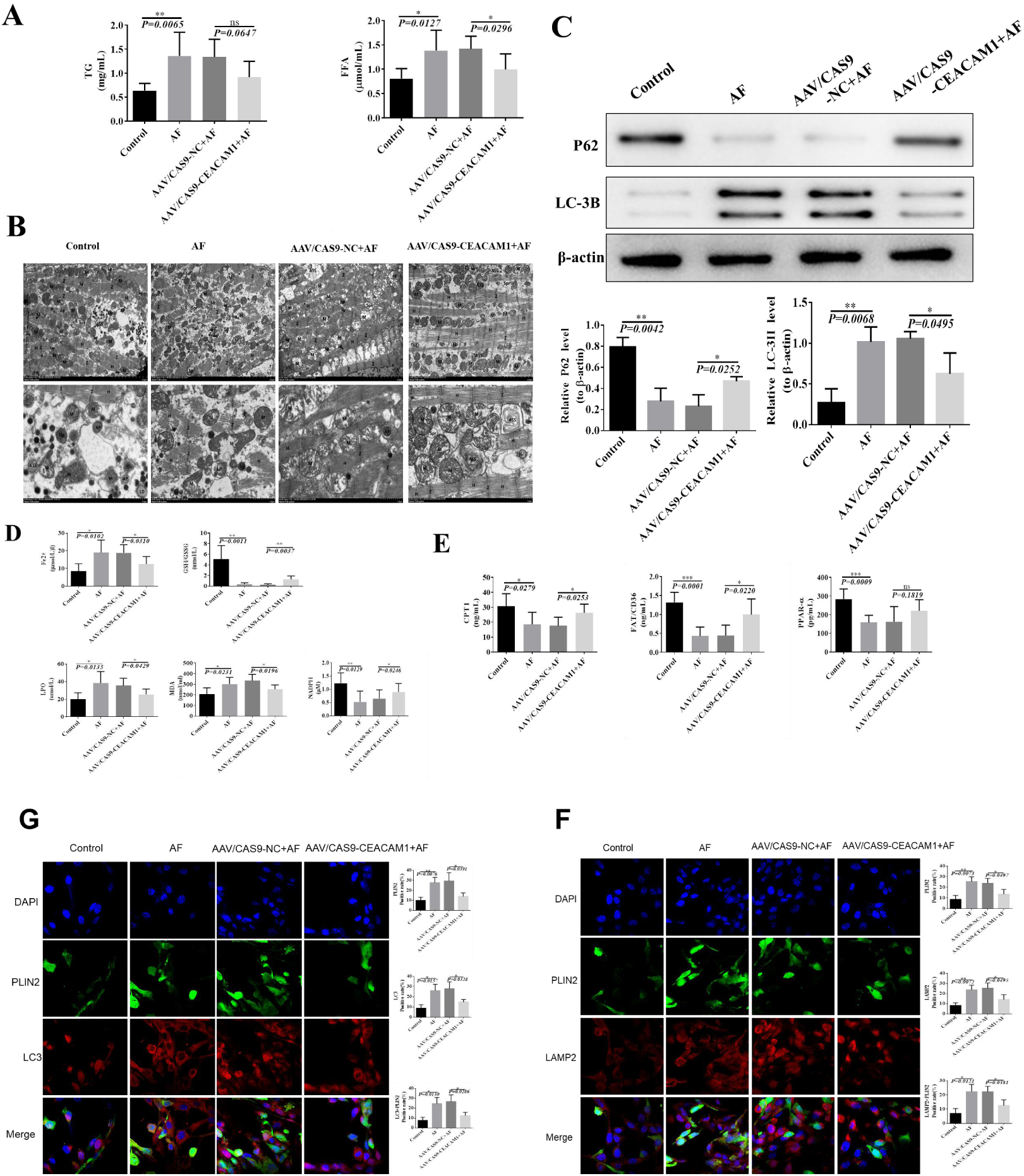
CEACAM1 promotes ferroptosis by inducing lipophagy to release FFA, elucidating its role in atrial fibrillation. A, TG and FFA content changes; B, Electron microscopy imaging of mitochondrial changes; C, Detection of autophagy-related proteins; D, Detection of ferroptosis-related factors; E, Key proteins involved in lipid metabolism; F, Co-localization of lipid droplets and lysosomes ; G, Co-localization of lipid droplets and autophagosomes.

## Disscussion

Ferroptosis is a recently identified form of regulated cell death distinct from apoptosis, characterized by iron-dependent lipid peroxidation. Although ferroptosis is closely associated with various cardiovascular diseases, its specific regulatory mechanisms in atrial fibrillation (AF) remain unclear. In this study, we demonstrated that CEACAM1 plays a critical role in lipophagy and ferroptosis by upregulating ABCA1 during the development of atiral fibrillation. This conclusion is supported by several in vivo and in vitro findings: (I) Ferroptosis is typically accompanied by lipid peroxide production, cellular iron accumulation, glutathione (GSH) depletion, and increased reactive oxygen species (ROS). Knockout of CEACAM1 altered these indicators, thereby inhibiting ferroptosis in AF, with ABCA1 identified as the downstream gene. (II) Lipophagy occurs in AF and is inhibited upon CEACAM1 knockout, accompanied by reduced free fatty acid (FFA) levels. Reactivation of ferroptosis by adding a lipophagy activator indicates that ferroptosis in AF is lipophagy-dependent and regulated by ABCA1, III) The restoration of lipophagy and ferroptosis upon FFA addition following CEACAM1 knockout suggests that FFA participates in the regulation of ferroptosis and CEACAM1 activity in atrial fibrillation.

Ferroptosis is regulated by various substances, and we found that CEACAM1 knockout inhibited ferroptosis in AF cells. The three primary mechanisms of ferroptosis include iron metabolism, amino acid metabolism, and lipid peroxidation. Iron imported into cells via transferrin enters as ferric ion (Fe3+), which is converted into ferrous ion (Fe2+) by ferric reductases within endosomes and transported to the cytosol by divalent metal transporter 1. The Fenton reaction facilitates the shuttling of Fe^2+^ to Fe^2+^, contributing to lipid peroxidation and ROS generation^19^. Various regulatory proteins and receptors influence ferroptosis by modulating lipid peroxidation and mitochondrial function through iron metabolism. For example, heat shock protein beta-1 (HSPB1) and CDGSH iron-sulfur domain 1 (CISD1) are negative regulators of ferroptotic cancer cell death^20, 21^. Cysteine oxidation and GSH depletion also play significant roles in ferroptosis^22^. Ferroptosis can be triggered by directly inhibiting GPX4 or suppressing GSH synthesis, an essential GPX4 cofactor. Extracellular cystine is exchanged in a 1:1 ratio for intracellular glutamate via the plasma membrane cystine/glutamate antiporter (system Xc^−^) and converted into cysteine, essential for GSH production. Reduced cystine levels lead to GSH depletion, GPX4 inactivation, and ferroptosis^23^. Fatty acid peroxidation is another key mechanism in ferroptosis. Enzymes such as acyl-CoA synthetase long-chain family member 4 (ACSL4), lysophosphatidylcholine acyltransferase 3 (LPCAT3), and lipoxygenases (LOXs) catalyze the formation and oxidation of polyunsaturated fatty acids (PUFAs), promoting ferroptosis^24-26^. The generation of ROS is one of the characteristics of ferroptosis.ROS generation is a hallmark of ferroptosis and occurs due to redox imbalances, leading to oxidative stress and cellular damage. ROS can be produced through the Fenton reaction or by mitochondria and NADPH oxidase (NOX) ^26, 27^. Oxidative stress plays a crucial role in AF pathogenesis. Increased ROS levels in AF have been linked to local (atrial myocytes) and systemic (serum) nicotinamide dinucleotide phosphate oxidase (NOX) activity^28^. Oxidative stress disrupts atrial gap junctions, causing electrical conduction abnormalities. For instance, excessive NOX2 activation reduces CX40 and CX43 expression, leading to atrial remodeling^28^. Oxidative stress also activates NF-κB, downregulating calcium channels and contributing to AF^29, 30^. Research has confirmed ferroptosis involvement in AF. Rapid pacing significantly reduces antioxidant gene expression in vivo and in vitro, while continuous ROS accumulation damages cell membranes through lipid peroxidation, leading to cell death^31^. Studies have also shown abnormal ferroptosis-related protein expression in AF, suggesting a link between ferroptosis and atrial fibrosis^19, 32^. CEACAM1, a member of the carcinoembryonic antigen (CEA) gene family, is known to regulate immune responses and metabolism. It has been implicated in conditions such as cancer and liver injury. Previous research indicates that CEACAM1 upregulation increases oxidative stress, affecting endothelial cell proliferation and causing cellular damage^22^. In this study, CEACAM1 knockout reduced oxidative stress in AF, characterized by decreased intracellular Fe^2+^ and Fe levels, increased NADPH and GSH/GSSG, and inhibited ferroptosis by reducing FFA. Considering the three major mechanisms of ferroptosis, CEACAM1 appears to be an indirect regulator of this process.

For the first time, we discovered that knocking out CEACAM1 can reduce free fatty acid (FFA) levels, inhibit lipophagy, and thereby suppress ferroptosis both in vivo and in vitro, suggesting a novel regulatory mechanism for ferroptosis. The synthesis, storage, and degradation of neutral lipids, such as triacylglycerols (TAG) and steryl esters, are dynamic processes^33^. Dysregulated lipid metabolism has been linked to inflammation, immunity, and cell death^34^. Proteomics and metabolomics studies have identified energy metabolism disorders in the myocardium of patients with AF, with lipid metabolism remodeling potentially contributing to AF development^35, 36^. Lipids are essential components of cell membranes, which contain high levels of polyunsaturated fatty acids (PUFAs) esterified into phospholipids and free cholesterol. These lipids help maintain cell structure and regulate function. However, the unstable C-H bonds in PUFAs are highly susceptible to attack by reactive oxygen species (ROS), resulting in lipid peroxidation^37^. Modulating the levels of enzymes involved in lipid metabolism can reduce PUFA peroxidation and mitigate ferroptosis caused by lipid toxicity. The degree of PUFA peroxidation is a critical determinant of ferroptosis susceptibility^38^. PUFAs oxidize to form lipid peroxidation products such as 4-hydroxynonenal (4-HNE) and malondialdehyde (MDA). These products, along with lipid peroxidation itself, destabilize and increase the permeability of cell membranes, ultimately leading to ferroptosis^39, 40^. Fatty acids can be stored in lipid droplets (LDs) as TAG and released through lipolysis when required^41^. The relationship between ferroptosis and lipid droplets is complex. During ferroptosis, lipophagy mediates the degradation of intracellular lipid droplets, releasing FFAs and decreasing lipid storage. Research has shown that FFAs released via lipophagy in liver cells provide substrates for lipid oxidation-dependent ferroptosis^42^. Consistent with our findings, CEACAM1 knockout and ABCA1 upregulation inhibited lipophagy, reduced FFA levels, and suppressed ferroptosis. However, other studies suggest that FFAs alone are not the driving factor for ferroptosis; instead, activated PUFAs must bind to membrane phospholipids (PLs) to induce peroxidation and exert toxic effects^43^. Phospholipids are key components of biomembranes, with their sn-2 positions typically esterified with acyl residues, including PUFAs. This attachment makes phospholipids highly susceptible to oxidation^44^. ABCA1 is a lipid transporter that mediates the transfer of cellular phospholipids and free cholesterol to extracellular matrix-associated apoA-I and related proteins^45^. Our study demonstrated that increased ABCA1 expression reduces ferroptosis, potentially by transporting phospholipids out of cells, thereby decreasing lipid peroxidation. However, conflicting findings have been reported in renal clear cell carcinoma, where Celastrol activates lipophagy via the LXRα/ABCA1 pathway, degrading lipid droplets and inhibiting cancer cell proliferation, migration, and invasion^46^. This contrasts with our observation that ABCA1 upregulation reduces lipophagy. Thus, further research is necessary to clarify the specific mechanisms by which ABCA1 inhibits lipophagy and ferroptosis.

Our study has certain limitations. For instance, monounsaturated fatty acids (MUFAs), another class of FFAs, can replace PUFAs in cell membranes, reducing the sensitivity of membrane lipids to oxidation ^47^. We did not investigate whether the FFAs increased by ABCA1 upregulation are predominantly PUFAs or MUFAs. Further studies are needed to elucidate the precise mechanisms underlying ABCA1-mediated inhibition of lipophagy and ferroptosis.

## Author Contributions

WW H. and JH Z. performed development of methodology and writing, AF model construction, review and revision of the paper; ZX L. and BN Y. performed acquisition, analysis and interpretation of data, and statistical analysis; X M., R Z. and J P. provided data collection and labelled the image, J W. and L Z. provided study concept and design, organization of experiment, as well as technical and material support. All authors read and approved the final pape.

